# Evidence for the role of music on the growth and signal response in duckweed

**DOI:** 10.1101/2022.03.25.485777

**Authors:** Zi Ye, Rui Yang, Ying Xue, Xinglin Chen, Ziyi Xu, Qiuting Ren, Jinge Sun, Xu Ma, Lin Yang

## Abstract

Sound vibration, an external mechanical force, has been proved to modulate plant growth and development like rain, wind, and vibration. However, the role of music on plants, especially on signal response, has been usually neglected in research. Herein, we investigated the growth state, gene expression, and signal response in duckweed treated with soft music. The protein content in duckweed after music treatment for 7 days was about 1.6 times that in duckweed without music treatment. Additionally, Fv/Fm in duckweed treated with music was 0.78, which was significantly higher in comparison with the control group (P < 0.01). Interestingly, music promoted the Glu and Ca signaling response. To further explore the global molecular mechanism, we performed transcriptome analysis. A total of 1296 DEGs were found for all these investigated genes in duckweed treated with music compared to the control group. Among these, up-regulation of the expression of metabolism-related genes related to glycolysis, cell wall biosynthesis, oxidative phosphorylation, and pentose phosphate pathways were found. Overall, these results provided a molecular basis to music-triggered signal response, transcriptomic, and growth changes in duckweed, which also highlighted the potential of music as an environmentally friendly stimulus to promote improved protein production in duckweed.

## Introduction

Sound is an oscillatory concussive pressure wave transmitted through gas, liquid, and solid. The influence of environmental factors such as water, wind, light, and predation on plant growth has been well studied ^[1]^. It has been shown that plants can perceive external sounds and some can even make sounds through the xylem ^[2]^ and communicate information through sound ^[3]^. Music affected the growth and physiological development of plants. The cabbage and cucumber at the seedling and maturity stages were treated with music, and it was found that oxygen uptake by the plants was significantly increased at both stages ^[4]^. Different types of sound enhanced plant survival ^[5]^, directed plant root growth ^[6]^, and shorted plant germination ^[7]^, thus sound influenced plant development. It has also been shown that certain sounds can affect fruit development. There were several examples including delaying fruit ripening ^[6,8]^ increasing the nutrient content of the fruit ^[9]^, and affecting flower pollination ^[10]^. However, further investigation addressing the mechanism of music influence on plants growth needs to be addressed.

It has been reported that sound also caused changes in the biochemical and genetic levels of plants. Sound has played a role in altering plant cell cycle, stomatal opening, growth hormone release, enzyme and hormone activity, immune function, RNA content, and transcript levels ^[1,11,12]^: (1) Sound altered the cell cycle of plants, accelerating protoplasmic movement within plant leaves ^[13]^ and speeding up plant metabolism. (2) Sound affected the stomatal opening of plant leaves ^[11]^, which led to faster uptake of nutrients and water ^[14]^. It has also allowed plants to absorb herbicides and pesticides, leading to the effective use of herbicides and pesticides. (3) The data suggested that some responses elicited by sound vibrations were modulated by specific changes in phytohormone levels, which also caused changes in phytohormone signaling like some stimuli ^[12]^. For example, the concentration of the salicylic acid increased in Arabidopsis under the influence of the sound of 1000 Hertz with 100 decibels, which increased plant defense and enhanced plant disease resistance ^[15]^. (4) In response to sound stimulation, some genes were activated at the transcriptional level ^[16]^, resulting in the changes in genes involved in the cellular metabolism of plants. Changes in gene expression and proteomics of Arabidopsis thaliana exposed to acoustic waves were studied. The expression of enzymes related to scavenging reactive oxygen species (ROS) and primary metabolism (tricarboxylic acid cycle, ATP synthesis, amino acid metabolism, and photosynthesis) has been upregulated ^[12]^. This showed that music had great potential for application in the field of plant biology. Therefore, it is important to figure out what molecular signaling processes activate those genes in plant growth during music environment.

Unlike animals with a rapid nervous system to responds to the environment, plants process several molecule signal transduction responses. Glutamate (Glu), an excitatory neurotransmitter in the mammalian central system, has been found to respond to the environment and play a role during plant growth. Glu has been proved to be a signaling molecule in plant cells and played an important role in seed germination and cellular metabolism ^[17–18]^. Glu, respond to mechanical damage, cold stress, and heat stress. ^[18–20]^ An interesting signal response caused by Glu was that it led to Calcium (Ca) signal responses throughout a whole plant body after wounding ^[19]^. This was due to Ca^2+^ flux depending on glutamate receptor-like proteins (GLR), Ca^2+^ permeable channels. Ca is one of the most versatile signals in living organisms ^[21]^, which play a significant role in the growth and development of plants in response to the environment: (1) Ca^2+^ was a key regulator, and its concentration in eukaryotic cells was significantly regulated by the Ca^2+^ sensor by the hormone ^[21–22]^; (2) Ca^2+^ binding protein signaling pathways mitigated biotic stresses ^[23]^; (3) Several tolerance strategies coordinated by Ca^2+^ signal has been found. For example, it responded to salt stress mediated by an enhanced antioxidant ^[24]^. These facts indicated that intracellular Ca^2+^ played an important role in eukaryotic signaling networks. However, it remains unknown that the response mechanisms of endogenous Glu and Ca^2+^ in duckweed under music treatment. The molecule signal responses in plants during music environment needs to be discovered.

Duckweed is a fast-growing aquatic plant, with high environmental adaptability and wide distribution ^[25]^. Compared to algae, duckweed is more readily available in large quantities, thus saving time and reducing energy consumption ^[26]^. The advantages of duckweed have high nutrient uptake and are commonly used for wastewater treatment to reduce environmental pollution ^[27]^. Furthermore, with its high protein and starch accumulation capacity, duckweed is widely used as an ideal raw material for animal feed and ethanol production ^[28]^. The potential of duckweed as an environmental indicator has also been studied and developed ^[29]^. However, due to the high protein content of duckweed, which is often used as the feed source, therefore it is especially significant to enhance the content of protein ^[30]^.

In this study, we used *Lemna turionifera 5511* as the material to study the change of duckweed treated with music. The main objectives were as follows: (i) to investigate the phenotypic and the content of protein in duckweed treated with music; (ii) to examine situations of Ca^2+^ signaling and Glu responses in duckweed treated with music; (iii) to reveal changes of significant metabolic pathways under music treatment; (iv) to study the gene expression changes and response mechanisms in duckweed under music treatment. The results of this study provided insight into how duckweed responded to music and how music can promote growth and the molecule mechanism of duckweed.

## Result

### Enhanced growth of duckweed by music

In order to investigate the physiological effects of duckweed treated with music, duckweed treated with or without music in 7 days has been studied. Shown as Fig. 1a and b, the frond number of duckweed distinctly increased after the 5th day of music treatment, although no significant change in phenotype was observed during the first 5 days of music treatment. The founds number of duckweed with music treatment for 6 d was 38, which is 3 more than that treated without music. Duckweed is a potential high-protein feed resource due to the protein content, thus the protein content of duckweeds with or without music was measured. Shown as Fig. 1c, the protein content in duckweeds treated with music after 7 days was significantly higher than the protein content in control duckweeds, which is 1.6 fold of that, These results showed that music in this study has positive effects on the growth of duckweed.

**Fig 1.**
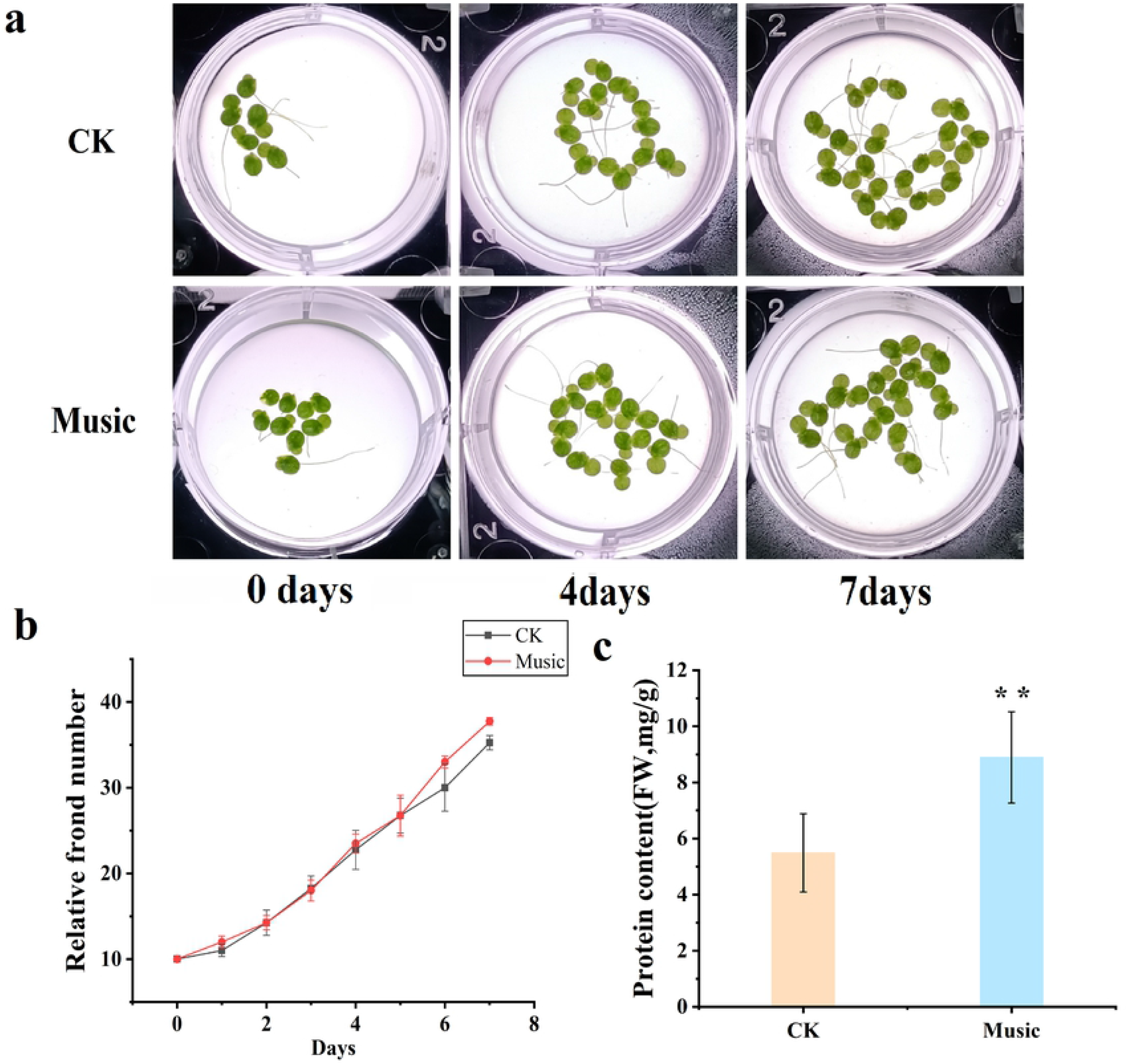
a. The phenotype of duckweed treated with or without music for 0-7 days; b. Frond growth curve over 7 days of cultivation treated with or without music; c. The contents of protein in duckwccd treated with or without music were determined by using Coomassic brilliant blue staining. The error bars in the graph are standard errors (SE) with biological repetitions n=5. *P<0.05, **P<0.01 represent significant differences and extremely significant differences.

### Enhanced photosynthetic capacity of duckweed under music treatment

The growth and nutrient accumulation of duckweed are closely related to photosynthetic capacity. The potential maximum photochemical efficiency of photosystem II (Fv/Fm), an intrinsic indicator of the photosynthetic capacity of plants, has been investigated in this study. As shown in Fig. 2a, Fv/Fm in duckweed treated with music was 0.78, which was significantly higher in comparison with the control group (P < 0.01). It was concluded that the photosynthetic capacity of the duckweed was enhanced under the condition of soothing music.

**Fig 2.**
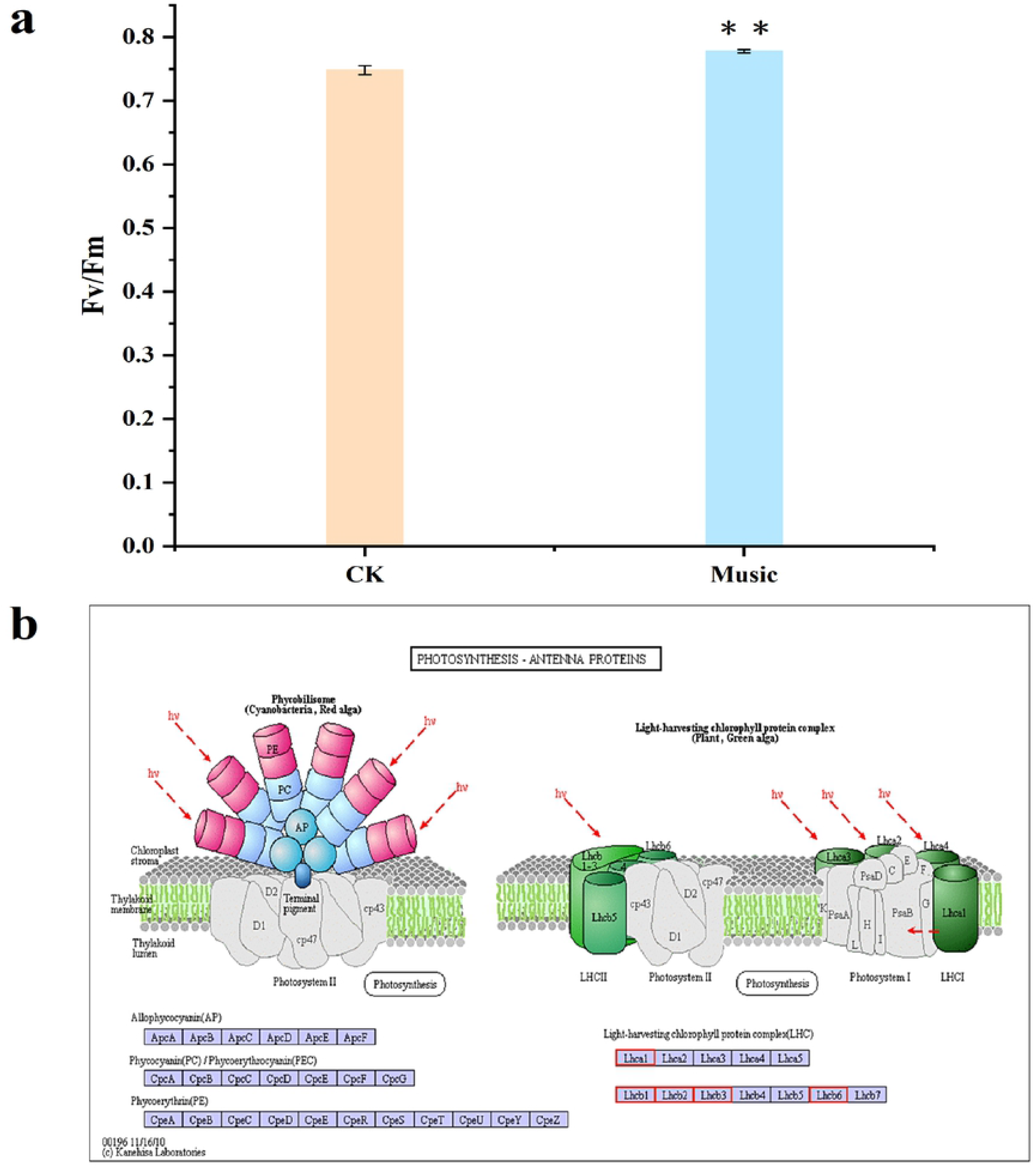
a. Fv/Fm was determined after treatment with and without music. The error bars in the graph arc standard errors (SE) with biological repetitions n=4.*P<0.05, **P<0.01 represent significant differences and extremely significant differences; b. Kyoto Encyclopedia of Genes and Gcnomcs (KEGG) maps the pathways associatcd with photosynthetic antenna proteins. The box color was determined by the expression pattern of unigcncs coding corresponding proteins. Red represents up-regulation, green represents down-regulation, and yellow represents mixed-regulation.

The gene expression difference in photosynthesis-antenna proteins-related gene expression has been investigated in the music group vs the control group (CK). Light-harvesting complex II (LHCII), the largest photosynthetic pigment-protein complex of plant photosystem II, has been up-regulated in the music group. Figure2b showed the pathway associated with photosynthesis-antenna proteins, and the red box represents up-regulated gene expression. Therefore, these results provided reasonable evidence for elevated photosynthesis during music treatment.

### Glu response and Glu contents under music treatment

Glu, playing a key role in protein composition, metabolism, and signaling has been investigated during music treatment. GFP-based Glu sensor iGluSnFR transgenic duckweed (iGlu duckweed) was successfully obtained in our previous study (Yang et al. Unpublished data). The fluorescence images of Glu in the roots of iGluSnFR duckweed subjected to transient music for 0-16 min were shown in Fig. 3a. The fluorescence intensity changed with music shock. The results indicated that music promoted the Glu signaling response of iGluSnFR duckweed.

**Fig 3.**
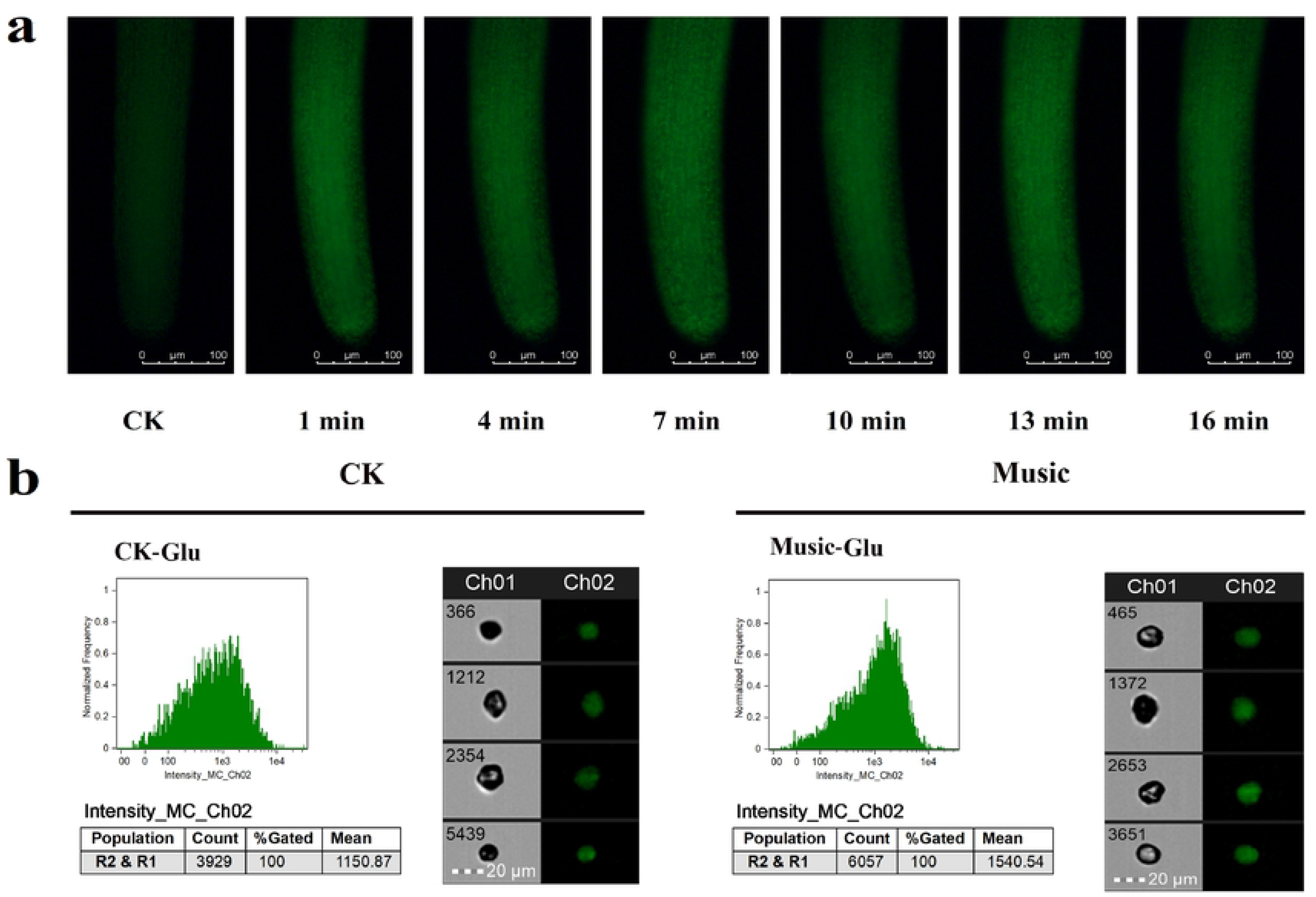
a. The fluorescence in the iGluSnFR duckweed subjected to transient music for 16 min; b. The Glu fluorescence intensity of the protoplasts in the roots of iGluSnFR duckweed was detected by flow cytometry, 6 thousand cells each group were analyzed. The iGluSnFR duckwccd was treated with or without music for 10 h. The excitation wavelength was 488nm. Scale bar=20 μ m(Ch 01 is the protoplast in bright, Ch 02 is fluorescently excited tong at 488 nm).

To further study the Glu signal response duckweed, the fluorescence intensity of protoplasts extracted from the iGluSnFR duckweed was analyzed under 488 nm excitation(Fig. 3b). The fluorescence intensity of iGluSnFR duckweed treated with or without music was 1540.54 and 1150.87, respectively. Thus, quantitative analysis showed that the fluorescence intensity of iGluSnFR duckweed subjected to soothing music was significantly stronger than those treated without music. The result revealed the involvement of the Glu signal during music processing.

### Differentially expressed genes (DEGs) and Gene ontology and KEGG pathway analyses of DEGs

The DEGs were analyzed and organized to better understand the molecular mechanisms of duckweed under music treatment. As shown in Fig. 4a, a total of 1296 DEGs were found in *“*Music vs CK*”*, of which 759 were down-regulated genes and 537 were up-regulated genes. In addition, the overall expression profile of the DEGs was observed in Fig. 4b, the heat map showed the gene expression levels that were significantly up-regulated and down-regulated in the case of *“*Music*”* and *“*CK*”*. The darker the red, the higher the expression. The darker the green, the lower the expression.

**Fig 4.**
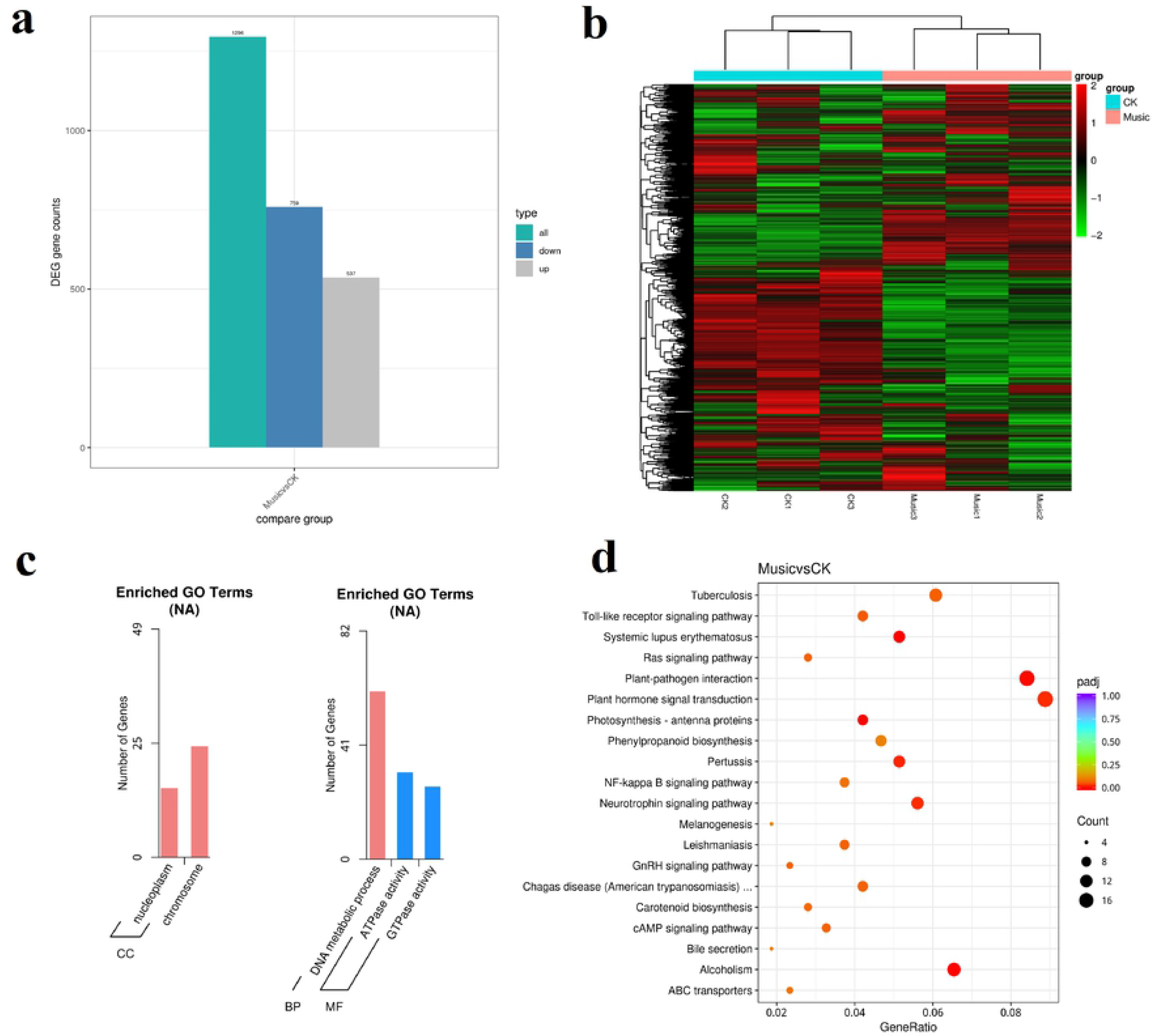
a. The histogram showed differentially expressed gcncs in response to music treatment; b. Cluster analysis of DGEs in the “Music” and “CK”; c. In the case of "Music vs CK”, the number of enriched up-regulated DEGs and down-regulated DEGs in different gene ontology categories. d. KEGG enrichment scatter plot (Rich factor referred to the ratio of the number of differentially expressed gcncs in the pathway to the total number of all annotated genes in the pathway).

To further understand the differences of DEGs in duckweed treated with music, GO enrichment analysis was performed on duckweed under music treatment and control group. As shown in Fig. 4c, *“*chromosome*”* was in the category of the cellular compartment with the most up-regulated DEGs. It was followed by *“*nucleoplasm*”* in the category of the cellular compartment. In contrast, the *“*DNA metabolic process*”* was in the category of the biological process with the most down-regulated DEGs. It was followed by *“*ATPase activity*”*, and *“*GTPase activity*”* in the category of molecular function.

Transcriptome analysis was performed to investigate the potential functions of KEGGs and DEGs in duckweed treated with music. The genes involved in plant hormone signal transduction and plant-pathogen interaction were significantly up-regulated and down-regulated, respectively. Among the 20 pathways with the most significant enrichment (Fig. 4d), the plant hormone signal transduction and plant-pathogen interaction contained the most differential genes.

### The Ca^2+^ signal response under music treatment

To further investigate the Ca signal response duckweed, the fluorescence intensity of protoplasts extracted from WT duckweed was analyzed under 488 nm excitation (Fig. 5). The average fluorescence intensity of 6000 protoplasts in WT duckweed treated with or without music were 1818.92 and 1582.06, respectively. Thus, quantitative analysis showed that the fluorescence intensity of WT duckweed subjected to soothing music was significantly stronger than those treated without music. The result revealed the involvement of the Ca signal during music processing.

**Fig 5.**
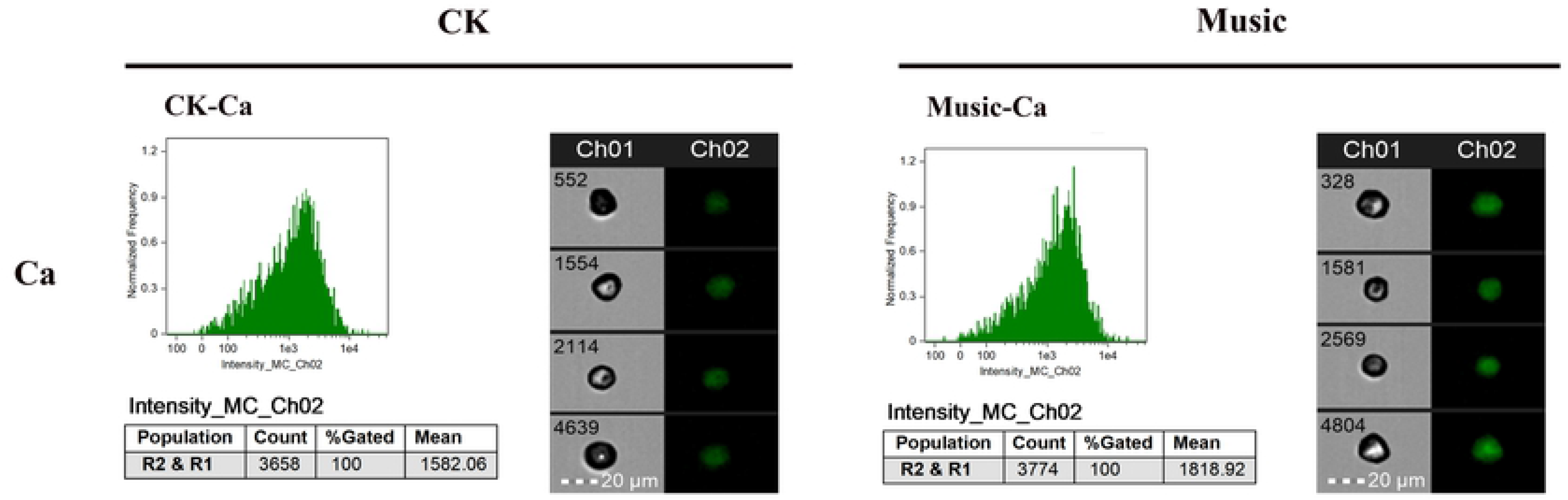
The duckweed was treated with or without music for 10 h and stained by Flou-4 AM. Ca content in protoplast analyzed by flowsight system in 488nm. Scale bar=20 μ m (Ch 01 is the protoplast in bright, Ch 02 is fluorescently excited tong at 488 nm).

To explore the differences in molecular mechanisms between duckweed treated with music and the control group, genes related to the Ca^2+^ signaling pathway were analyzed. As shown in Fig. 6, GTP-binding protein (GPCR) was up-regulated, which has raised 2.17 log2 Fold Change. While adiponectin receptor (AdipoR) located on the plasma membrane was mix-regulated. Moreover, calmodulin (CAM) genes, Calcium-dependent protein kinase (CPK) genes, and AMP-activated protein kinase (AMPK) genes were down-regulated. Furthermore, calcineurin B-like protein (CBL) has raised 1.04 log2 Fold Change. And Ras-related C3 botulinum toxin substrate 1 (Rac) genes were up-regulated, which has raised 5.43 log2 Fold Change. While CREB-binding protein (CBP) genes were down-regulated. These results suggested that mTOR, CAMK, CBL, Rac, CBP, and RBOH were triggered by intracellular calcium ions, and the expression of proteins related to metabolism has also altered.

**Fig 6.**
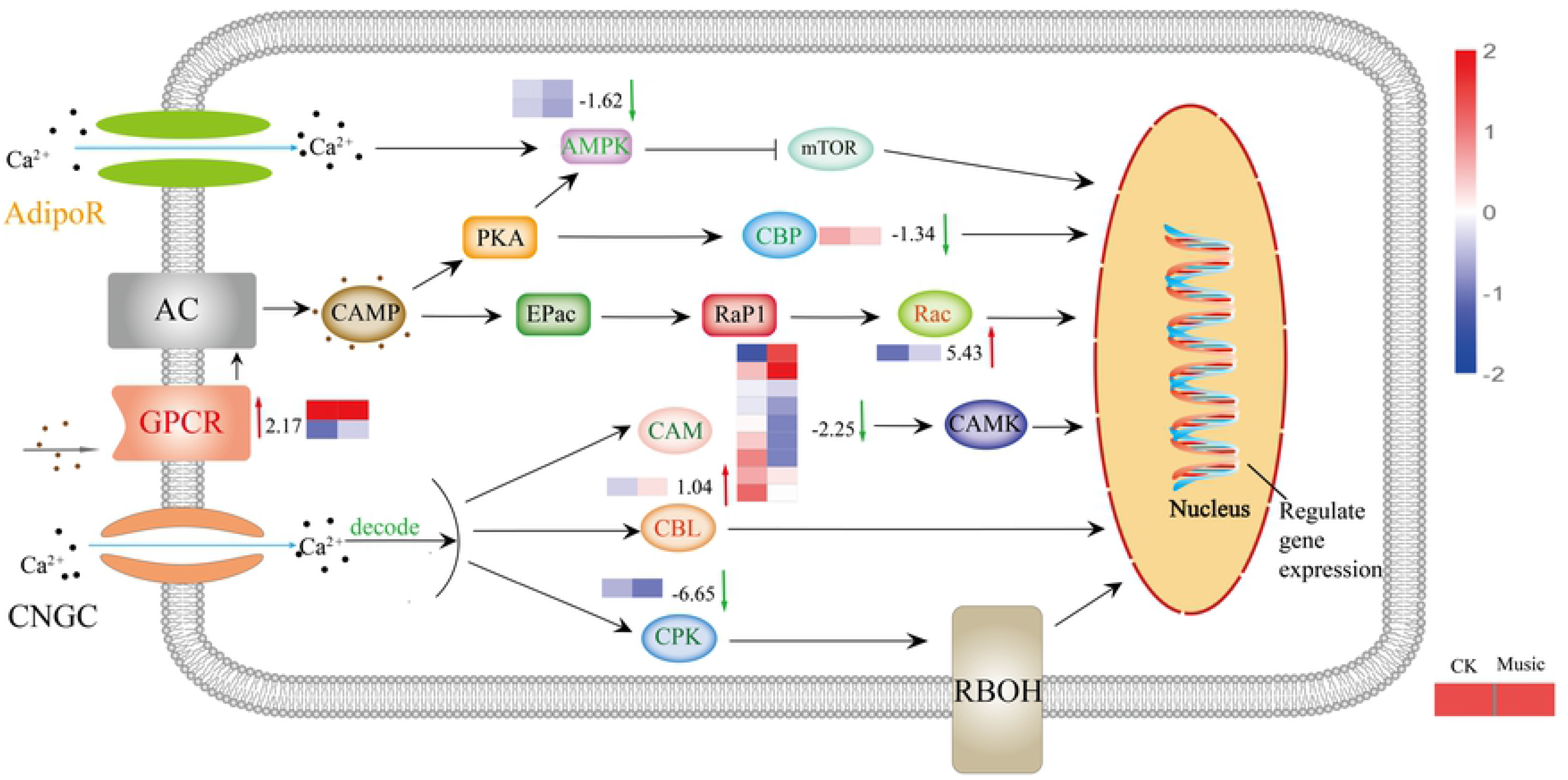
The response of the Ca signal pathway under music treatment. The channels and proteins associated with Ca signal transduction were shown in the figure. The color of letters and arrows represented the change in the DEGs: red indicated up-regulated, green indicated down-regulated, and yellow indicated mix-regulated. The heat map next to the arrow showed the expression of different genes encoded the protein in Music and CK duckweed. Red meant high expression; blue meant low expression. (Rae: Ras-related C3 botulinum toxin substrate 1, AdipoR: adiponcctin receptor, CPK: crcatinc kinasc, AMPK: AMP-activated protein kinasc, GPCR: GTP-binding protein, CBP: CREB-binding protein, CBL: calcincurin protein, CAM: calmodulin)

### Metabolic pathways of duckweed under music treatment

Some primary and secondary metabolic pathways related to the molecular mechanism were represented in Fig. 7. Starch is considered to be an energy deposit in plants. The gene encoding glucose-1-phosphate adenyltransferase (G1PA) is up-regulated during starch metabolism, which has raised 1.00 log2 Fold Change. However, the gene encoding alpha-amylase (AA) is down-regulated. GSH, with the help of glutathione S-transferase (GST), is transformed into R-S-glutathione. The glutathione S-transferase (GST) gene in glutathione metabolism was up-regulated, which has raised 1.56 log2 Fold Change. Further, the gene encoded 6-phosphogluconate dehydrogenase (PGD) was up-regulated involved in the pentose phosphate pathway, which has raised 1.52 log2 Fold Change. Therefore, it tended to be activated to provide more NADPH under music treatment. The expression of pyruvate kinase (PK) taking part in glycolysis was down-regulated. Additionally, NADH-ubiquinone oxidoreductase chain 5 (ND5) mix-regulated and NAD(P)H-quinone oxidoreductase subunit 2 (NdhB) was up-regulated in the oxidative phosphorylation pathway were also noteworthy, which has raised 1.17 log2 Fold Change. In addition, the biosynthesis of pectin is one of the major branches of sugar metabolic flux, and most of the enzymes in this pathway were up-regulated in duckweed.

**Fig 7.**
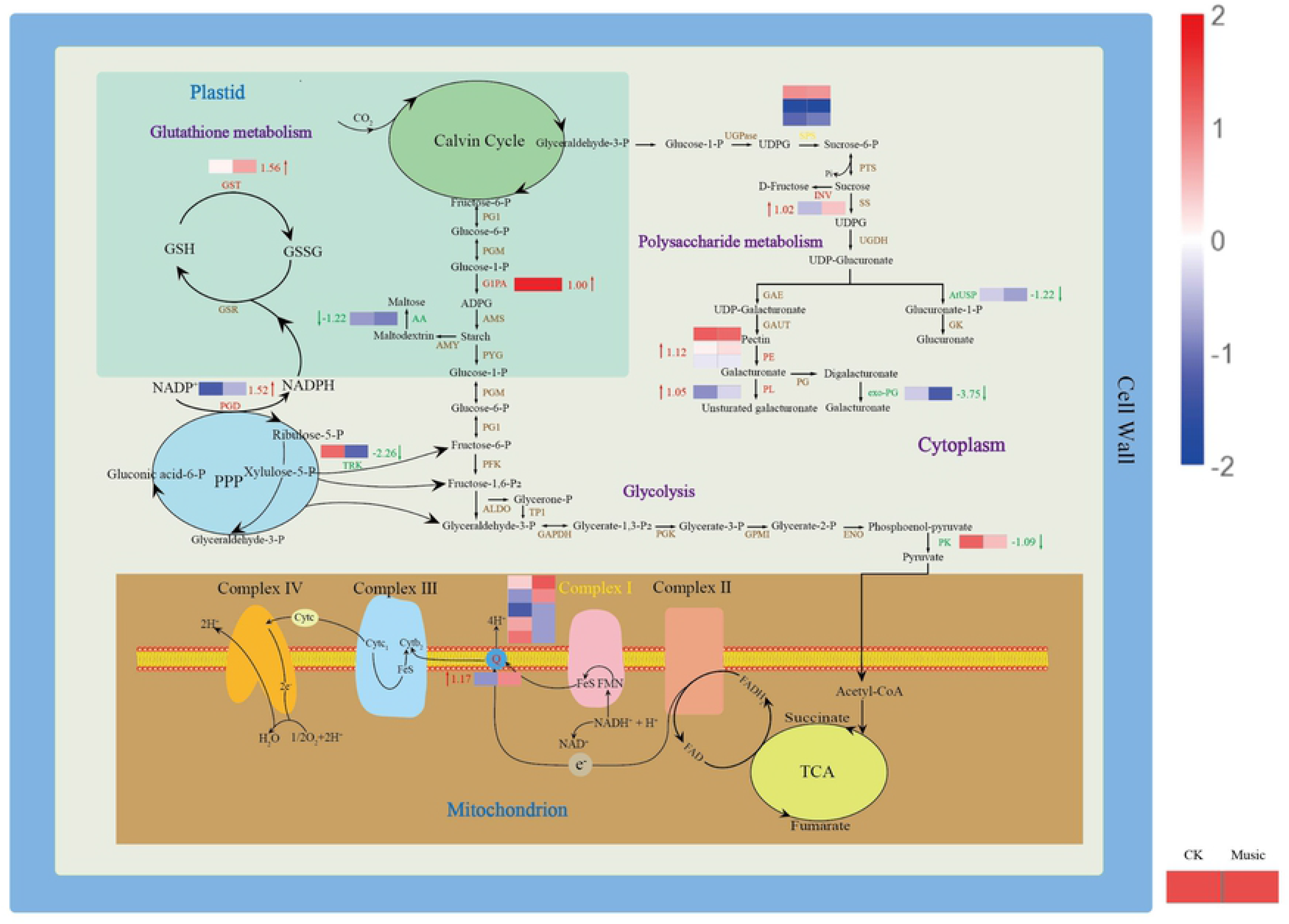
The metabolic mechanism of duckweed was influenced by music. Metabolites (in black) and proteins with enzymes and transporters are shown. The arrows beside or through proteins indicate the directions of catalytic reactions or transportation. Rcd arrows represent uprcgulation, grccn arrows represent downregulation, and yellow arrows represent mixed regulation. The heat map next to the arrow showed the expression of different genes encoded the protein in Music and CK duckweed. Red meant high expression; blue meant low expression. (INV: in+vertase, SPS: sucrose-phosphate synthasc, G 1 PA: glucose-l-phosphatc adenylyltransferase, PK: pyruvate kinasc, PE: pcctincstcrasc, PL: pcctatc lyasc, cxo-PG: galacturan 1,4-alpha-galacturonidasc, AtUSP: UDP-sugar pyrophosphorylasc, AA: alpha-amylase, GST: glutathionc S-transfcrasc, PGD:6-phosphogluconatc dchydrogcnasc, TRK: transkctolasc, ND5: NADH-ubiquinone oxidorcductasc chain 5, NdhB: NAD (P) H-quinone oxidorcductasc subunit 2)

## Discussion

The growing price of Soybean meal presents a challenge to researchers to investigate the possibilities for applicating green technologies to increase the protein production of plants to replace soybean as the feed resource. The Sound caused changes in the growth and development, biochemical and genetic levels of plants ^[1,4]^. Researchers found that sound waves significantly affected the number of leaves of mustard greens ^[14]^. In our preliminary research, we found that different decibels have different effects on the growth of duckweed. Music at too high a decibel level was not beneficial to plants. This was consistent with the study of Wang et al. and Cai et al. ^[7,31]^ In this study, the results showed that the soft music named “The Purple Butterfly” at 60-70 decibels promoted the growth of the frond number of duckweed (Fig. 1). Furthermore, under music treatment for 7d, the average protein content in duckweed treated with music was 8.89 mg/g FW, which was significantly higher than the average protein content (5.49 mg/g FW) in duckweed treated without music (Fig. 1c). A similar result has been reported that sound waves enhanced the synthesis of soluble protein in chrysanthemum ^[16]^. Therefore, music promoted the accumulation of protein in duckweed.

Nucleic acid, the hereditary material, stores a digital record of every protein’s design. The transcript is important for protein expression. It has been reported that the transcript level in HL-60 cells under the electric magnetic field was enhanced by four times compared to the control ^[32]^. Besides, environmental factors such as water stress led to an obvious effect on the synthesis of d protein. In our research, we studied the effects of soft music on transcription. GO enrichment analysis has been performed and it showed that ‘chromosome’ was in the category of the cellular compartment with the most up-regulated DEGs (Fig.4c). Thus, the enhancement of protein accumulation by music might be related to the DEGs associated with chromosome synthesis being up-regulated. The findings demonstrated that music increased the utilization of duckweed in terms of its use as the feed source.

Fv/Fm reflects the maximum light energy conversion efficiency of the PSⅡreaction center. Meng found that Fv/Fm was increased of strawberry leaves under the sound wave ^[33]^, which was similar to our research. Fv/Fm in duckweed treated with music was higher than that of CK (Fig. 2a), which suggested that music enhanced the activity of the PS II reaction center and the ability to use light energy. In addition, most of the DEGs related to ‘Photosynthesis-antenna proteins’ were up-regulated (Fig. 2b). This result suggested that antenna proteins genes were closely associated with increased photosynthesis under music. Notably, in a previous study, Lhcb6 has been shown to be involved in reducing oxidative stress and photoprotection under natural conditions ^[35]^. In addition to this, it has been found that Lhcb2 was associated with a light-harvesting complex playing a critical role in providing the energy required for photolysis ^[12]^. Therefore, music stimulation promoted photosynthesis by increasing the expression of photosynthesis-related genes with music treatment.

The understanding of the genome of duckweed has greatly facilitated the application of phylogenetic studies^[34]^. And genome of duckweed provided a great opportunity to investigate how different levels of metabolism translate into molecular changes of genes ^[36]^. In the present research, functional annotation of DEGs via the database of KEGG revealed that “Plant hormone signal transduction” was highly enriched in the music group compared with control (Fig. 4d). A similar result has been reported that most of the genes involved in hormonal signaling were also up-regulated in arabidopsis treated with sound ^[12]^. Nevertheless, how music affected the expression of genes related to hormone signaling in duckweed was needed to further study.

Plants could sense local signals like wounding, and transmit the signal to the whole plant body rapidly by Glu and Ca^2+^ signal ^[19]^. Our results showed that music stimulated the Glu signaling response by iGluSnFR duckweed (Fig. 3a). We investigated changes in the Glu content of duckweed treated with music. The results showed that music enhanced the content of the signaling molecule Glu (Fig. 3b). Exactly, the Ca^2+^ signal, which could be released by GLR with Glu activation, acts as a second messenger and regulator in most signaling networks. The localization of Ca^2+^ stained by Fluorescence-4 AM showed that music increased the amount of Ca^2+^ accumulation (Fig. 5). Toyota et al. (2018) showed that mechanical stimulation elicited a change in plant Glu response, which led to a change in Ca^2+^ concentration to induce a local plant response. We noticed significant up-regulation of the CBL gene in duckweed, calcineurin B-like protein (CBL), which interacted with CBL-interacting protein kinases (CIPKs) to facilitate downstream signaling. Functions as a key component in the regulation of various stimuli or signals in plants^[37]^. Our results suggested that the second messenger Ca^2+^ may be involved in the signaling of musical stimuli. Among them, GTP-binding protein (GPCR) was also significantly up-regulated, affecting the influx of calcium ions and promoting signal transmission. Ghosh et al. (2016) have also shown that activation of ion transport proteins played a role in sound-mediated responses. In addition, calmodulin (CAM) and calcium-dependent protein kinase (CPK) were significantly down-regulated (Fig. 6), which played regulatory roles in a variety of cellular processes, affecting plant stress resistance and growth and development ^[38–39]^. However, the mechanisms underlying the regulation of resistance in duckweed under the influence of music still need to be further explored.

This study provided the first comprehensive transcriptome analysis of *Lemna turionifera 5511* exposed to music. Transcriptome analysis identified several biological processes involved in response to musical stimuli. Researchers have found that stimulation by external factors may lead to changes in the proteome associated with primary metabolism in plants ^[40]^. This was similar to the results of our study. These results showed that genes involved in the basic biological processes of carbohydrate metabolism and protein metabolism respond to musical stimuli, with most of them being up-regulated (Fig. 7). For example, the transcript levels of some genes involved in cellular in the pentose phosphate pathway and cell wall biosynthesis were up-regulated. Previous studies have shown that sound increased the content of soluble sugars in chrysanthemums ^[16]^. In the present experiment, genes involved in starch and sucrose metabolism were up-regulated, also suggesting that music promoted carbohydrate accumulation in plants. ATP, a commonly used carrier of energy in cells, is produced through two main pathways: light reactions in chloroplasts and oxidative phosphorylation in mitochondria. The transcript levels of the genes encoding the enzymes involved in oxidative phosphorylation were up-regulated in this study, suggesting that ATP content may be enhanced in response to musical stimulation during electron transport in the respiratory chain. This may be one of the reasons for the accelerated metabolic response. Our study identified changes in the metabolic pathways of *Lemna turionifera 5511* in response to musical stimulation through transcriptome analysis. The use of music to stimulate the expression of plant genes and promote plant growth and metabolism offers new ideas for the development of ecological agriculture.

In conclusion, we explored frond number, protein content, photosynthetic capacity, Glu and Ca^2+^ signaling response, genes analysis, and metabolic pathway analysis during music environment. Our data has allowed us to confirm music enhanced growth, especially protein content, by enhancing the expression of photosynthesis-related genes and stimulating the Glu signal response of duckweed. Delving into the fields of plant production and protein enhancement, we gained insight into the function of music. These results provided new ideas for research in the field of plant acoustics.

## Materials and methods

### Duckweed cultivation and music processing

The duckweeds (*Lemna turionifera 5511*) were cultured in our lab in a sterile environment following Yang et al (2013). The duckweed was treated with or without the music named “The Purple Butterfly” for 7 days. Music was played for five hours a day, with an intensity of 60-70 decibels. The culture conditions were with a light intensity of 62 μmol m^−2^ s^−1^ and a temperature of 20/28*°* C. The number of leaves was counted daily during the experiment, while the leaf area was calculated using Image J (NIH, Bethesda, MD, USA) software.

### Measurement of the photosynthetic system

Photosynthetic fluorescence parameters were measured in the duckweeds after music treatment. The samples were treated in dark for 30 min to deplete the organic matter in the leaves before measurement. Subsequently, the maximum quantum yield (Fv/Fm) of photosystem II (PSII) of duckweed leaves was measured by Dual-PAM100 fluorometer (Waltz Company, Germany).

### Determination of protein content

The duckweeds (0.5 g) treated with or without music were ground with 5 ml 0.05 M phosphate-buffered saline (PBS) and the protein was extracted. After adding Coomassie Brillant Blue G-250 staining solution to the standards and samples, they were mixed for 5 min. A series of gradients of standard protein solutions were prepared using standard bovine serum proteins. The absorbance of the standards and samples at 595 nm was recorded using an Enzyme-labeled Instrument. A standard curve was generated by plotting the average absorbance at 595 nm as a function of the concentration of the protein standard. The sample protein content can be calculated from the standard curve.

### Music promotes the fluorescent response of duckweed

The iGluSnFR duckweed was treated with music 10h in advance. And then we carried out experimental observations in a musical setting. Fluorescence changes in the roots of the duckweed under the influence of transient music were observed using a fluorescence microscope (Leica, DFC450C, MD5000, Berlin, Germany).

### Flow cytometric analysis of Ca and Glu content in duckweed

Protoplasts extraction: duckweed was treated with the music for 10 h. The duckweed was fixed in 95% ethanol for 15 min, and then washed 2-3 times by adding DPBS. The roots and leaves were separated and enzymatically digested by adding 1% pectinase and 1% cellulase at 37*°*C for 60 min in dark. The obtained protoplasts were washed three times with DPBS and then filtered through a 200-mesh cell sieve.

Glu content in protoplasts of duckweed analysis: The protoplasts of iGluSnFR duckweed treated with or without music were analyzed by FlowSight (Merck millipore, FlowSight® imaging flow cytometer, Germany). The exciting wavelength was 488 nm. 6000 protoplasts in each group were analyzed and 6 Six parallel groups were made. Ca^2+^ content in protoplasts of duckweed analysis: The WT duckweed was stained with Fluorescence-4 AM dye at 37° C in the dark for 1h. The protoplasts of WT duckweed treated with or without music were analyzed by FlowSight (Merck millipore, FlowSight® imaging flow cytometer, Germany). The exciting wavelength was 488 nm. 6000 protoplasts in each group were analyzed and 6 parallel groups were made.

### RNA library sequencing

The duckweed treated with or without music was frozen in a −80°C refrigerator and then sequenced at Novogene (Chaoyang, Beijing). Total RNA was extracted using the RNAprep Pure Plant Kit (TIANGEN, Beijing, China). RNA integrity was precisely checked using an Agilent 2100 bioanalyzer system and purity was checked using a NanoPhotometer® spectrophotometer (IMPLEN, CA, USA). Sequencing libraries were generated using the NEBNext® Ultra *™* RNA Library Prep Kit (NEB, USA). After library construction, initial quantification was performed using a Qubit 2.0 Fluorometer to dilute the library to 1.5 ng/ul, followed by detection of the insert size of the library using an Agilent 2100 bioanalyzer system. After meeting expectations, qRT-PCR was performed to accurately quantify the effective concentration of the library (effective library concentration higher than 2nM) to ensure the quality of the library. After passing the library test, Illumina NovaSeq 6000 (Illumina, USA) sequencing was performed at Novogene Bioinformatics Technology Co., Ltd., Beijing.

### Differential expression analysis and functional annotation

Reference sequences were compared using RSEM ^[41]^ (bowtie2 default parameters) software, and these reads were converted using FPKM ^[42]^, the most commonly used method for estimating gene expression levels. The resulting reads were the input data for differentially expressed genes (DEGs). For samples with biological duplicates, we used DESeq2^[43]^, with differential gene screening criteria of |log2(FoldChange)| > 1 & padj < 0.05.

Gene function annotations were selected from the following authoritative databases: protein sequence database (Swiss-Prot), Gene Ontology (GO), Clusters of Orthologous Groups of proteins (KOG/COG), Protein family (Pfam), NCBI non-redundant nucleotide sequences (Nt), KEGG Ortholog database (KO) and NCBI non-redundant protein sequences (Nr).

### KEGG and GO enrichment analysis

The GO enrichment analysis method was followed by Young et al (2010), which was based on Wallenius non-central hyper-geometric distribution. It provided a more accurate calculation of the probability of GO term enrichment by differential genes, with GO enrichment being considered significant with padj less than 0.05. KOBAS software was used to perform KEGG functional enrichment analysis on the differential gene set. Pathway significant enrichment analysis was performed using KEGG Pathway as the unit, Hypergeometric test was applied to find out the pathway where the differential gene was significantly enriched relative to all annotated genes. KEGG pathway enrichment was likewise used with padj less than 0.05 as the threshold for significant enrichment.

### Data Screening and Analysis

Data from the results of at least three independent parallel experiments were taken, organized, and calculated using Microsoft Excel 2010 and then analyzed using SPSS software (IBM SPSS Statistics, Version 20), p-values were calculated using independent samples t-test, p < 0.05 indicates significant, p < 0.01 indicates highly significant, and finally Plotting was performed using the software ORIGIN 2018 (asterisks indicate significant differences: *P < 0.05, **P < 0.01).

## Acknowledgments

We would like to express our special thanks of gratitude to soft music entitled “The Purple Butterfly” by Bandari for its contribution to our research.

## Supporting information

**S1 File. The music score for this research.** The file is the score of the soft music used for this research entitled “The Purple Butterfly” by Bandari.

## Author Contributions

**Data curation:** Ying Xue, Rui Yang
**Formal analysis:** Lin Yang
**Investigation:** Rui Yang, Xinglin Chen, Jinge Sun
**Methodology:** Ying Xue, Ziyi Xu, Qiuting Ren
**Project administration:** Zi Ye, Lin Yang
**Resources:** Zi Ye, Lin Yang
**Software:** Rui Yang, Ying Xue, Xinglin Chen
**Supervision:** Lin Yang, Zi Ye
**Validation:** Lin Yang, Xu Ma
**Visualization:** Rui Yang, Ying Xue, Ziyi Xu
**Writing – original draft:** Lin Yang, Ying Xue, Rui Yang
**Writing – review & editing:** Lin Yang, Rui Yang, Ying Xue

## Data availability statement

### Competing financial interests

The authors declare no competing financial interests. All relevant data are within the manuscript and its Supporting Information files.

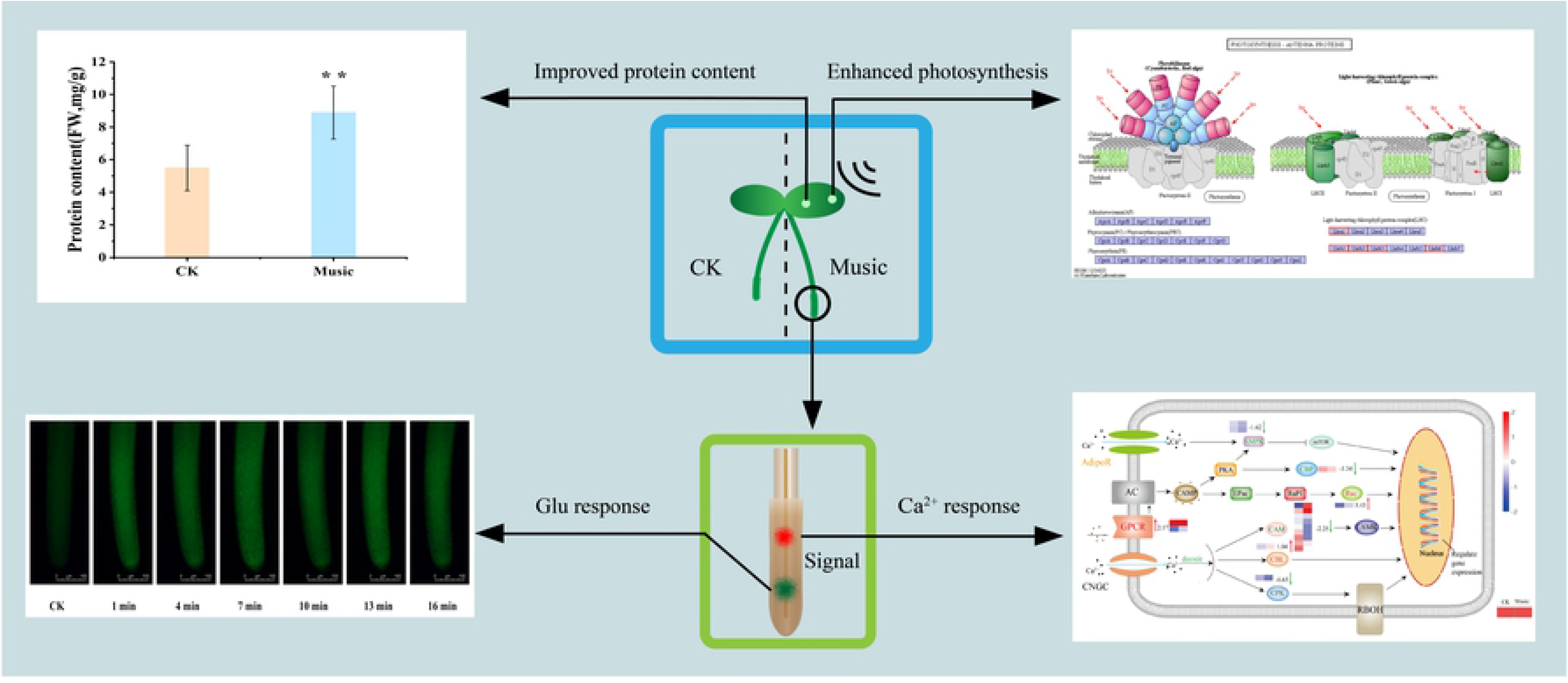

## References

1. Reda HE Hassanien, Tian-zhen HOU, Yu-feng LI, Bao-ming LI. Advances in Effects of Sound Waves on Plants. Journal of Integrative Agriculture. 2014; 13(2): 335–348. https://dx.doi.org/10.1016/S2095-3119(13)60492-X.

2. Laschimke R, Burger M, Vallen H. Acoustic emission analysis and experiments with physical model systems reveal a peculiar nature of the xylem tension. J Plant Physiol. 2006 Oct;163(10):996–1007. https://doi.org/10.1016/j.jplph.2006.05.004. PMID: 16872717.

3. Gagliano M. Green symphonies: a call for studies on acoustic communication in plants. Behav Ecol. 2013 Jul;24(4):789–796.https://doi.org/10.1093/beheco/ars206.PMID: 23754865.

4. Qin YC, Lee WC, Choi YC, Kim TW. Biochemical and physiological changes in plants as a result of different sonic exposures. Ultrasonics. 2003 Jul;41(5):407–11. https://doi.org/10.1016/s0041-624x(03)00103-3. PMID: 12788223.

5. López-Ribera I, Vicient CM. Drought tolerance induced by sound in Arabidopsis plants. Plant Signal Behav. 2017 Oct 3;12(10):e1368938. https://doi.org/10.1080/15592324.2017.1368938. PMID: 28829683

6. Kim JY, Lee HJ, Kim JA, Jeong MJ. Sound Waves Promote Arabidopsis thaliana Root Growth by Regulating Root Phytohormone Content. Int J Mol Sci. 2021 May 27;22(11):5739. https://doi.org/10.3390/ijms22115739. PMID: 34072151

7. Cai W, He H, Zhu S, Wang N. Biological effect of audible sound control on mung bean (Vigna radiate) sprout. Biomed Res Int. 2014;2014:931740. https://doi.org/10.1155/2014/931740. PMID: 25170517

8. Joo-Yeol Kim, Jin-Su Lee, Taek-Ryoun Kwon, Soo-In Lee, Jin-A. Kim, Gyu-Myoung Lee, et al. Sound waves delay tomato fruit ripening by negatively regulating ethylene biosynthesis and signaling genes. Postharvest Biology and Technology. 2015;110: 43–50. https://doi.org/10.1016/j.postharvbio.2015.07.015

9. Altuntas O, Ozkurt H. The assessment of tomato fruit quality parameters under different sound waves. J Food Sci Technol. 2019 Apr;56(4):2186–2194. https://doi.org/10.1007/s13197-019-03701-0. PMID: 30996452

10. Veits M, Khait I, Obolski U, Zinger E, Boonman A, Goldshtein A, et al. Flowers respond to pollinator sound within minutes by increasing nectar sugar concentration. Ecol Lett. 2019 Sep;22(9):1483–1492. https://doi.org/0.1111/ele.13331. PMID: 31286633

11. Rajagopalan UM, Wakumoto R, Endo D, Hirai M, Kono T, Gonome H, et al. Demonstration of laser biospeckle method for speedy in vivo evaluation of plant-sound interactions with arugula. PLoS One. 2021 Oct 28;16(10):e0258973. https://doi.org/10.1371/journal.pone.0258973. PMID: 34710145

12. Ghosh R, Mishra RC, Choi B, Kwon YS, Bae DW, Park SC, et al. Exposure to Sound Vibrations Lead to Transcriptomic, Proteomic and Hormonal Changes in Arabidopsis. Sci Rep. 2016 Sep 26;6:33370. https://doi.org/10.1038/srep33370. PMID: 27665921

13. Frongia F, Forti L, Arru L. Sound perception and its effects in plants and algae. Plant Signal Behav. 2020 Dec 1;15(12):1828674. https://doi.org/10.1080/15592324.2020.1828674. PMID: 33048612

14. Y Hendrawan, Hendrawan Y, Anniza K N, Prasetyo J, Djoyowasito G. Effect of plant sound wave technology to increase productivity of mustard greens (Brassica juncea L.). IOP Conference Series: Earth and Environmental Science. 2020; 524(1): 012012.

15. Choi B, Ghosh R, Gururani MA, Shanmugam G, Jeon J, Kim J, et al. Positive regulatory role of sound vibration treatment in Arabidopsis thaliana against Botrytis cinerea infection. Sci Rep. 2017 May 30;7(1):2527. https://doi.org/10.1038/s41598-017-02556-9. PMID: 28559545

16. Wang X J, Wang B C, Jia Y, Duan C R, Sakanishi A. Effect of sound wave on the synthesis of nucleic acid and protein in chrysanthemum. Colloids and Surfaces B: Biointerfaces. 2003;29:99–102. https://doi.org/10.1016/S0927-7765(02)00152-2

17. Kong D, Ju C, Parihar A, Kim S, Cho D, Kwak JM. Arabidopsis glutamate receptor homolog3.5 modulates cytosolic Ca2+ level to counteract effect of abscisic acid in seed germination. Plant Physiol. 2015 Apr;167(4):1630–42. https://doi.org/10.1104/pp.114.251298. PMID: 25681329

18. Li ZG, Ye XY, Qiu XM. Glutamate signaling enhances the heat tolerance of maize seedlings by plant glutamate receptor-like channels-mediated calcium signaling. Protoplasma. 2019 Jul;256(4):1165–1169. https://doi.org/10.1007/s00709-019-01351-9. PMID: 30675652.

19. Toyota M, Spencer D, Sawai-Toyota S, Jiaqi W, Zhang T, Koo AJ, et al. Glutamate triggers long-distance, calcium-based plant defense signaling. Science. 2018 Sep 14;361(6407):1112–1115. https://doi.org/10.1126/science.aat7744. PMID: 30213912.

20. Zheng Y, Luo L, Wei J, Chen Q, Yang Y, Hu X, Kong X. The glutamate receptors AtGLR1.2 and AtGLR1.3 increase cold tolerance by regulating jasmonate signaling in Arabidopsis thaliana. Biochem Biophys Res Commun. 2018 Dec 2;506(4):895–900. https://doi.org/10.1016/j.bbrc.2018.10.153. Erratum in: Biochem Biophys Res Commun. 2021 Aug 20;566:211-213. PMID: 30392908.

21. Liang J, He Y, Zhang Q, Wang W, Zhang Z. Plasma Membrane Ca2+ Permeable Mechanosensitive Channel OsDMT1 Is Involved in Regulation of Plant Architecture and Ion Homeostasis in Rice. Int J Mol Sci. 2020 Feb 7;21(3):1097. https://doi.org/10.3390/ijms21031097. PMID: 32046032

22. Liu Q, Ding Y, Shi Y, Ma L, Wang Y, Song C, et al. The calcium transporter ANNEXIN1 mediates cold-induced calcium signaling and freezing tolerance in plants. EMBO J. 2021 Jan 15;40(2):e104559. https://doi.org/10.15252/embj.2020104559. PMID: 33372703

23. Patra N, Hariharan S, Gain H, Maiti MK, Das A, Banerjee J. TypiCal but DeliCate Ca++re: Dissecting the Essence of Calcium Signaling Network as a Robust Response Coordinator of Versatile Abiotic and Biotic Stimuli in Plants. Front Plant Sci. 2021 Nov 25;12:752246. https://doi.org/10.3389/fpls.2021.752246. PMID: 34899779

24. Seifikalhor M, Aliniaeifard S, Shomali A, Azad N, Hassani B, Lastochkina O, et al. Calcium signaling and salt tolerance are diversely entwined in plants. Plant Signal Behav. 2019;14(11):1665455. https://doi.org/10.1080/15592324.2019.1665455. PMID: 31564206

25. Yang GL, Feng D, Liu YT, Lv SM, Zheng MM, Tan AJ. Research Progress of a Potential Bioreactor: Duckweed. Biomolecules. 2021 Jan 13;11(1):93. https://doi.org/10.3390/biom11010093. PMID: 33450858

26. Sun Z, Guo W, Yang J, Zhao X, Chen Y, Yao L, Hou H. Enhanced biomass production and pollutant removal by duckweed in mixotrophic conditions. Bioresour Technol. 2020 Dec;317:124029. https://doi.org/10.1016/j.biortech.2020.124029. PMID: 32916457

27. Wang W, Messing J. Status of duckweed genomics and transcriptomics. Plant Biol (Stuttg). 2015 Jan;17 Suppl 1:10–5. https://doi.org/10.1111/plb.12201. PMID: 24995947

28. Yu C, Zhao X, Qi G, Bai Z, Wang Y, Wang S, et al. Integrated analysis of transcriptome and metabolites reveals an essential role of metabolic flux in starch accumulation under nitrogen starvation in duckweed. Biotechnol Biofuels. 2017 Jun 26;10:167. https://doi.org/10.1186/s13068-017-0851-8. PMID: 28670341

29. Stomp AM. The duckweeds: a valuable plant for biomanufacturing. Biotechnol Annu Rev. 2005;11:69–99. https://doi.org/10.1016/S1387-2656(05)11002-3. PMID: 16216774

30. Roman B, Brennan RA, Lambert JD. Duckweed protein supports the growth and organ development of mice: A feeding study comparison to conventional casein protein. J Food Sci. 2021 Mar;86(3):1097–1104. https://doi.org/10.1111/1750-3841.15635. PMID: 33624354.

31. Wang Bochu, Chen Xin, Wang Zhen, Fu Qizhong, Zhou Hao, Ran Liang. Biological effect of sound field stimulation on paddy rice seeds. Colloids and Surfaces B: Biointerfaces.2003; 32(1): 29–34. https://doi.org/10.1016/S0927-7765(03)00128-0.

32. Reba Goodman, Ann Shirley-Henderson. Transcription and translation in cells exposed to extremely low frequency electromagnetic fields, Journal of Electroanalytical Chemistry and Interfacial Electrochemistry.1991; 320(3): 335–355. https://doi.org/10.1016/0022-0728(91)85651-5.

33. Meng QW, Zhou Q, Zheng SJ, Gao Y. Responses on photosynthesis and variable chlorophyll fluorescence of Fragaria ananassa under sound wave. Energy Procedia.2012;16: 346–352.

34. Wang W, Haberer G, Gundlach H, Gläßer C, Nussbaumer T, Luo MC, et al. The Spirodela polyrhiza genome reveals insights into its neotenous reduction fast growth and aquatic lifestyle. Nat Commun. 2014;5:3311. https://doi.org/10.1038/ncomms4311. PMID: 24548928

35. Chen YE, Ma J, Wu N, Su YQ, Zhang ZW, Yuan M, et al. The roles of Arabidopsis proteins of Lhcb4, Lhcb5 and Lhcb6 in oxidative stress under natural light conditions. Plant Physiol Biochem. 2018 Sep;130:267–276. https://doi.org/10.1016/j.plaphy.2018.07.014. PMID: 30032070

36. Zhang Y, An D, Li C, Zhao Z, Wang W. The complete chloroplast genome of greater duckweed (Spirodela polyrhiza 7498) using PacBio long reads: insights into the chloroplast evolution and transcription regulation. BMC Genomics. 2020 Jan 28;21(1):76. https://doi.org/10.1186/s12864-020-6499-y. PMID: 31992185

37. Ma X, Li QH, Yu YN, Qiao YM, Haq SU, Gong ZH. The CBL-CIPK Pathway in Plant Response to Stress Signals. Int J Mol Sci. 2020 Aug 7;21(16):5668. https://doi.org/10.3390/ijms21165668. PMID: 32784662

38. Luan S, Kudla J, Rodriguez-Concepcion M, Yalovsky S, Gruissem W. Calmodulins and calcineurin B-like proteins: calcium sensors for specific signal response coupling in plants. Plant Cell. 2002;14 Suppl(Suppl):S389–400. https://doi.org/10.1105/tpc.001115. PMID: 12045290

39. Atif RM, Shahid L, Waqas M, Ali B, Rashid MAR, Azeem F, et al. Insights on Calcium-Dependent Protein Kinases (CPKs) Signaling for Abiotic Stress Tolerance in Plants. Int J Mol Sci. 2019 Oct 24;20(21):5298. https://doi.org/10.3390/ijms20215298. PMID: 31653073

40. Kosová K, Vítámvás P, Prášil IT, Renaut J. Plant proteome changes under abiotic stress--contribution of proteomics studies to understanding plant stress response. J Proteomics. 2011 Aug 12;74(8):1301–22. https://doi.org/10.1016/j.jprot.2011.02.006. PMID: 21329772.

41. Li B, Dewey CN. RSEM: accurate transcript quantification from RNA-Seq data with or without a reference genome. BMC Bioinformatics. 2011 Aug 4;12:323. https://doi.org/10.1186/1471-2105-12-323. PMID: 21816040

42. Trapnell C, Williams BA, Pertea G, Mortazavi A, Kwan G, van Baren MJ, et al. Transcript assembly and quantification by RNA-Seq reveals unannotated transcripts and isoform switching during cell differentiation. Nat Biotechnol. 2010 May;28(5):511–5. https://doi.org/10.1038/nbt.1621. PMID: 20436464

43. Love MI, Huber W, Anders S. Moderated estimation of fold change and dispersion for RNA-seq data with DESeq2. Genome Biol. 2014;15(12):550. https://doi.org/10.1186/s13059-014-0550-8. PMID: 25516281

44. Yang L, Han H, Liu M, Zuo Z, Zhou K, Lü J, et al. Overexpression of the Arabidopsis photorespiratory pathway gene, serine: Glyoxylate aminotransferase (AtAGT1), leads to salt stress tolerance in transgenic duckweed (Lemna minor). Plant Cell Tissue and Organ Culture 2013; 113(3): 407–416

45. Young MD, Wakefield MJ, Smyth GK, Oshlack A. Gene ontology analysis for RNA-seq: accounting for selection bias. Genome Biol. 2010;11(2): R14. https://doi.org/10.1186/gb-2010-11-2-r14. PMID: 20132535

